# A novel human monoclonal antibody specific to the A33 glycoprotein recognizes colorectal cancer and inhibits metastasis

**DOI:** 10.1101/748962

**Authors:** Patrizia Murer, Louis Plüss, Dario Neri

## Abstract

Colorectal cancer represents the second most common cause of cancer-related death. The human A33 transmembrane glycoprotein is a validated tumor-associated antigen, expressed in 95% of primary and metastatic colorectal cancers. Using phage display technology, we generated a human monoclonal antibody (termed A2) specific to A33 and we compared its epitope and performance to those of previously described clinical-stage anti-human A33 antibodies. All antibodies recognized a similar immunodominant epitope, located in the V-domain of A33, as revealed by SPOT-analysis. The A2 antibody homogenously stained samples of poorly, moderately and well-differentiated colon adenocarcinomas. All antibodies also exhibited an intense staining of healthy human colon sections. The A2 antibody, reformatted in murine IgG2a format, preferentially localized to A33-transfected CT26 murine colon adenocarcinomas in immunocompetent mice with a homogenous distribution within the tumor mass, while other antibodies exhibited a patchy uptake in neoplastic lesions. A2 efficiently induced killing of A33-expressing cells through antibody-dependent cell-mediated cytotoxicity *in vitro* and was able to inhibit the growth of A33-positive murine CT26 and C51 lung metastases *in vivo*. Anti-A33 antibodies may thus represent useful vehicles for the selective delivery of bioactive payloads to colorectal cancer, or may be used in IgG format in settings of minimal residual disease.

## INTRODUCTION

Colorectal cancer (CRC) is one of the most common cancer types worldwide and ranks second in terms of cancer-related deaths [1]. Approximately 20% of newly diagnosed CRC patients have already developed metastases in the liver, lungs, lymph nodes, peritoneum or soft tissues [2]. Despite significant progresses in standard care of therapies, including antibody-based therapeutics inhibiting angiogenesis (anti-VEGF-A) or tumor growth factors (anti-EGFR), the prognosis of patients with metastatic CRC remains poor [3]. Immune checkpoint inhibitors, which provide a clinical benefit for patients with many different types of malignancies, are typically not active for the treatment of colorectal cancer [4], exception made for DNA Mismatch Repair-Deficient or Microsatellite Instability-High metastatic CRC patients, which respond positively to the combination treatment of nivolumab (anti-PD-1) and ipilimumab (anti-CTLA-4) [5]. For these reasons, there is an urgent need to develop more efficacious strategies for the treatment of patients with metastatic CRC.

Intense research efforts are currently being devoted to the implementation of antibody-based therapeutic strategies, in which the antibody moiety serves as delivery vehicle for bioactive agents. Radionuclides [6], immunostimulatory cytokines [7, 8] and T cell engagers (e.g., bispecific antibodies; [9]) have been considered as payloads for the treatment of metastatic CRC. In this context, the carcinoembryonic antigen (CEA) and transmembrane glycoprotein A33 represent the most commonly used target antigens, which have received extensive Nuclear Medicine validation in patients, using radiolabeled preparations of monoclonal antibody reagents [6, 10–13].

A33 is expressed in epithelia of the lower gastrointestinal tract and in 95% of primary and metastatic colorectal cancers [14]. Clinical-stage antibodies have been developed against A33 and have been used for CRC treatment either as “naked” immunoglobulins [15], engineered for the delivery of beta-emitting radionuclides [6] or of T cell-engaging antibody moieties [16]. The first anti-A33 antibody to be used for tumor-targeting application was developed by the group of Lloyd J. Old [17]. In this article, it will be referred to as “K”. A humanized version of K (here described as K.hu) was reported in 1995 [18]. Recently, MacroGenics has described the isolation and characterization of a novel humanized monoclonal antibody (here described as “MG”), which has been used for the development of bispecific products in DART^®^ format [16].

Here we describe the isolation and characterization of a novel human antibody (termed A2), specific to A33. A2, which was compared to the K, K.hu and MG antibodies in terms of biochemical properties and tumor-targeting characteristics, was found to avidly bind to CRC specimens *in vitro* and *in vivo*. The novel anti-A33 antibody was able to kill A33-expressing murine adenocarcinoma cells by ADCC *in vitro* and prevent formation of murine CRC lung metastases *in* vivo. The results described in this article suggest that A2 may be applied as intact antibody able to kill colorectal tumor cells through ADCC, or as building block for the implementation of antibody-based therapeutic strategies for the treatment of metastatic CRC.

## RESULTS

### Murine model of colorectal cancer expressing human A33

The human transmembrane glycoprotein A33 is expressed in 95% of human colorectal cancers but has only 67% amino acid identity with its murine counterpart. In order to establish an immunocompetent murine model of colorectal cancer expressing A33, we stably-transfected the murine colorectal carcinoma cell lines CT26^wt^ and C51^wt^ with the gene coding for A33 [Figure 1a]. The resulting clones CT26^A33.C3^ and C51^A33.A5^, selected by antibiotic resistance and single-cell sorting, showed a shift in fluorescence intensity upon FACS analysis using A33-specific antibodies, compared to isotype controls [Figure 1a]. The staining intensities were comparable with the one observed using the LS174T cells human colorectal cancer cell line.

**Figure 1.**
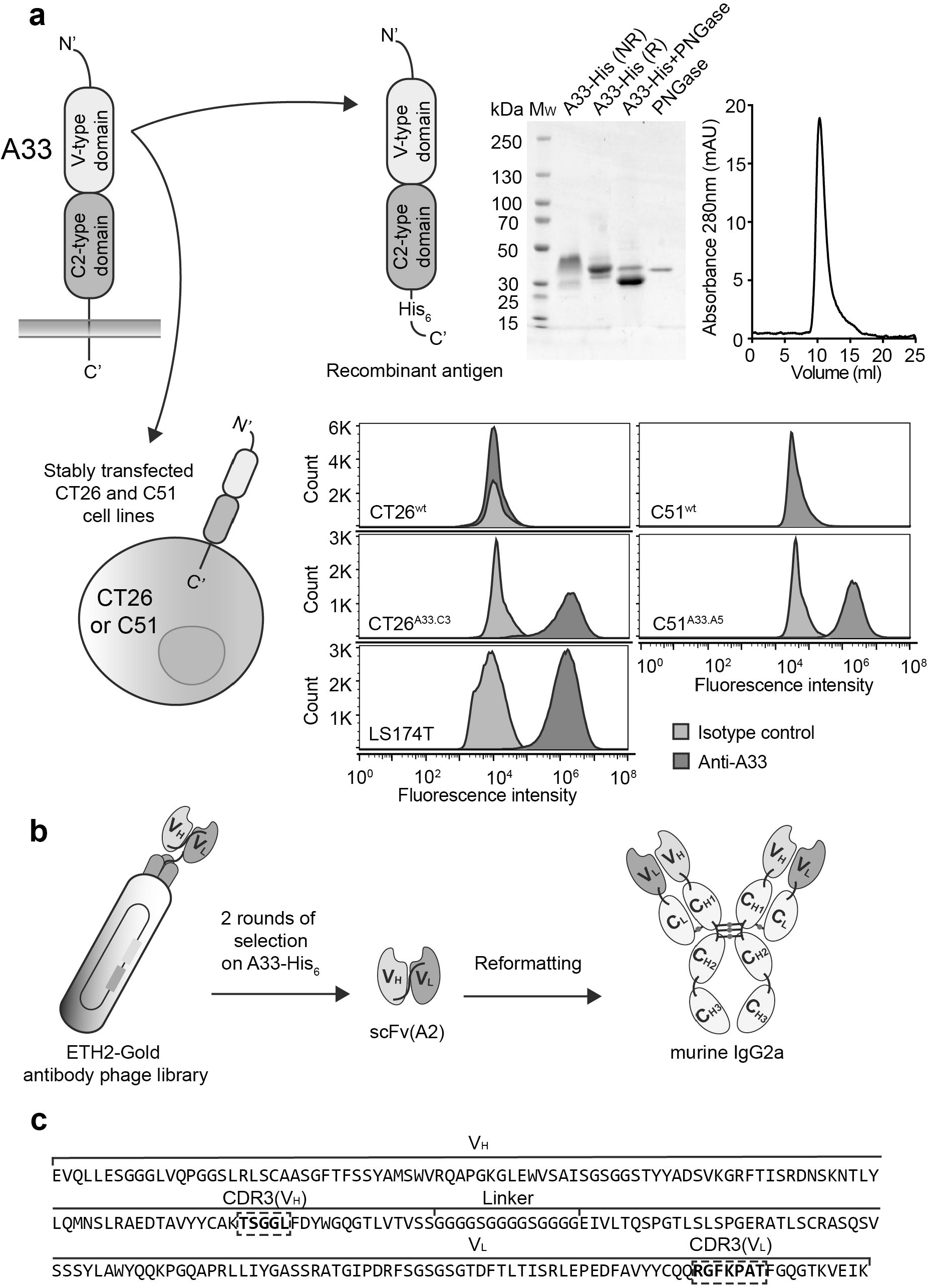
Recombinant A33-His antigen, murine A33-expressing adenocarcinoma cell lines and novel anti-A33 antibody. **a) Top left:** Schematic representation of the transmembrane A33 antigen. **Top right:** Schematic representation of the recombinant A33-His protein expressed in CHO cells. Only the extracellular V-type and C2-type Ig-like domains were expressed with a C-terminal Histidine Tag (His_6_). The recombinant antigen was characterized by SDS-PAGE analysis (MW: molecular weight, NR: non-reducing, R: reducing) and size exclusion chromatography. **Bottom left:** Representative image of murine colorectal cell lines CT26 and C51 expressing the A33 antigen as transmembrane protein. **Bottom right:** FACS analysis detecting expression of A33 on CT26^wt^, C51^wt^, human colorectal tumor cell line LS174T and transfected CT26^A33.C3^ and C51^A33.A5^ cells. Staining was performed with a murine anti-A33 specific antibody and the corresponding signal was amplified with an anti-mouse AlexaFluor488 secondary antibody. An isotype-matched antibody was used as negative control. **b)** Schematic representation of the selection of the A33 specific antibody A2, as scFv fragment, from the ETH-2-Gold phage display library. The scFv(A2) was reformatted into murine IgG2a for further characterization. **c)** Amino acidic sequence of scFv(A2) as selected from the ETH-2-Gold library, with the portions of the CDR3 loops of the VH and VL domains highlighted by dashed squares.

### Expression of recombinant A33-His and antibody isolation

The extracellular Ig-like domains of A33 were cloned with a His-tag at the C-terminus into the mammalian expression vector pcDNA3.1(+) for protein expression in CHO cells [Figure 1a]. The purified recombinant glycoprotein A33-His eluted as a single peak in size exclusion chromatography and migrated as a large band in non-reducing SDS-PAGE analysis, revealing a higher molecular weight (Mw) than the calculated Mw of 24.4 kDa. The sequence of A33 [**Supplementary Figure 1**] contains three possible sites for N-linked glycosylation. Deglycosylation of A33-His with PNGase led to the formation of a smaller and uniform band, with apparent Mw of 30kDa in SDS-PAGE analysis [Figure 1a].

### Isolation of an A33 specific monoclonal antibody by phage display

The human antibody A2 in scFv format was isolated by panning the ETH-2-Gold phage display library [19] against A33-His, immobilized on a solid support. ScFv(A2) was reformatted into a chimeric murine IgG2a format for mammalian cell expression [Figure 1b]. Figure 1c indicates the amino acid sequence of scFv(A2), highlighting the portions of the CDR3 loops of V_H_ and V_L_ domains that had been combinatorially mutated in the ETH-2-Gold library.

### Expression and characterization of anti-A33 antibodies in the IgG2a format

In order to compare the antibody A2 to previously described clinical-stage anti-A33 antibodies [K [17], K.hu [18] and MG [9]], all antibodies were reformatted in murine IgG2a format. The proteins were expressed in CHO cells and purified to homogeneity, using Protein A chromatography. The amino acid sequences of K, K.hu and MG are reported in **Supplementary Figure 2**. Figure 2 illustrates the biochemical properties and the BIAcore profiles of the four antibodies. A slow dissociation from the antigen was observed for the A2 and MG antibodies, while a biphasic dissociation profile was seen for K and K.hu (the latter antibody showing a more rapid dissociation from A33 immobilized on the BIAcore sensor chip) [Figure 3]. Analyses of the functional affinities on CT26^A33.C3^ cells with serial dilutions of the anti-A33 IgG2a antibodies revealed apparent K_D_ values ranging from 2-20pM [Figure 4a]. The stronger functional affinity of K and K.hu to A33 on CT26^A33.C3^ cells compared to A2 and MG, does not match with the BIAcore profiles on the immobilized A33. TA99, specific to a melanoma antigen [20], was used as negative control of irrelevant specificity in this context.

**Figure 2.**
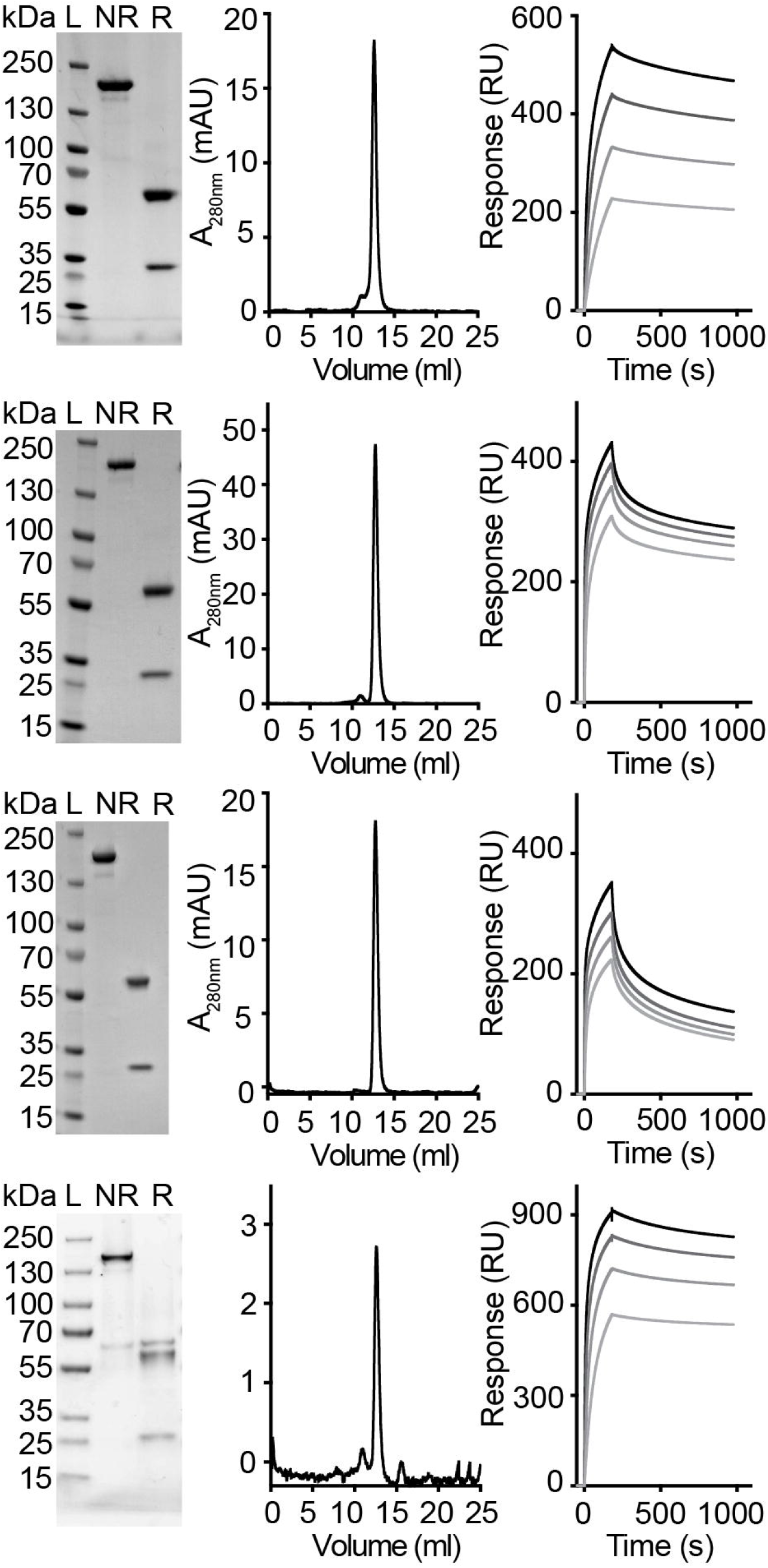
*In vitro* characterization of anti-A33 antibodies in the murine IgG2a format. SDS-PAGE analyses under non-reducing (NR) and reducing (R) conditions, size exclusion chromatography profiles and SPR analyses of IgG2a(A2), IgG2a(K), IgG2a(K.hu) and IgG2a(MG). SPR analyses performed at antibody concentrations of 63nM, 125nM, 250nM and 500nM.

**Figure 3.**
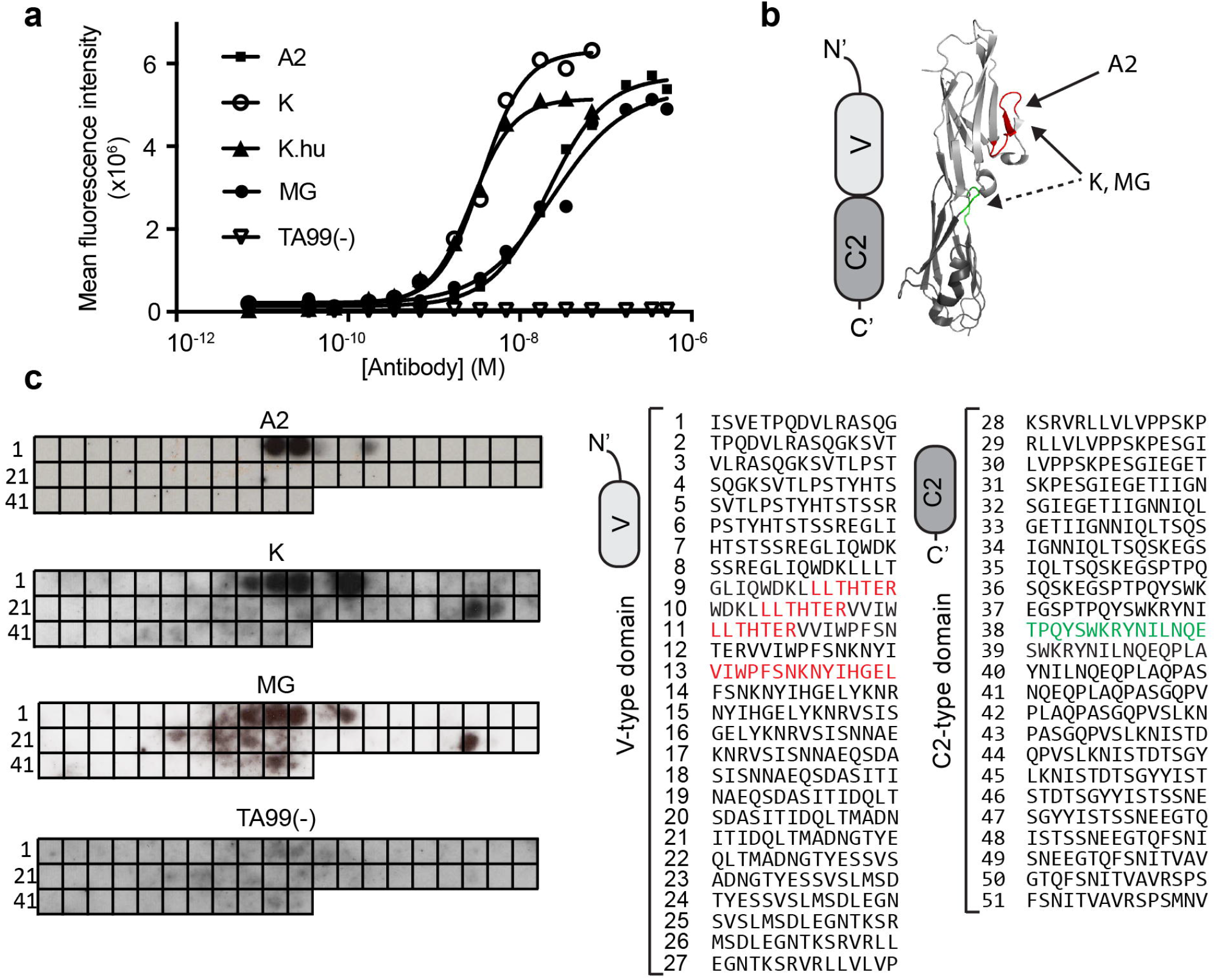
Binding affinity of the anti-A33 antibodies on CT26^A33.C3^ and epitope mapping. **a)** Functional binding affinity of the antibodies A2, K, K.hu, MG and negative control TA99(-), measured by flow cytometry on CT26^A33.C3^ cells. **b)** Schematic representation and crystal structure of V-type and C2-type Ig-like domains, indicating the binding epitopes of A2 (red), K and MG (discontinuous epitope red + green). **c. Left:** Binding epitopes of the anti-A33 antibodies A2, K and MG and the negative control TA99(-) on the PepSpot membrane (as black spots). Each spot (1-51) on the membrane is coated with 15 amino acid-long peptides, spanning the amino acid sequence of A33. **Right:** Amino acidic sequence of every spot, with binding epitopes on the Ig-like V-type domain (for all three anti-A33 antibodies) in red and binding epitopes on the Ig-like C2-type domain (for the antibodies K and MG) in green.

**Figure 4.**
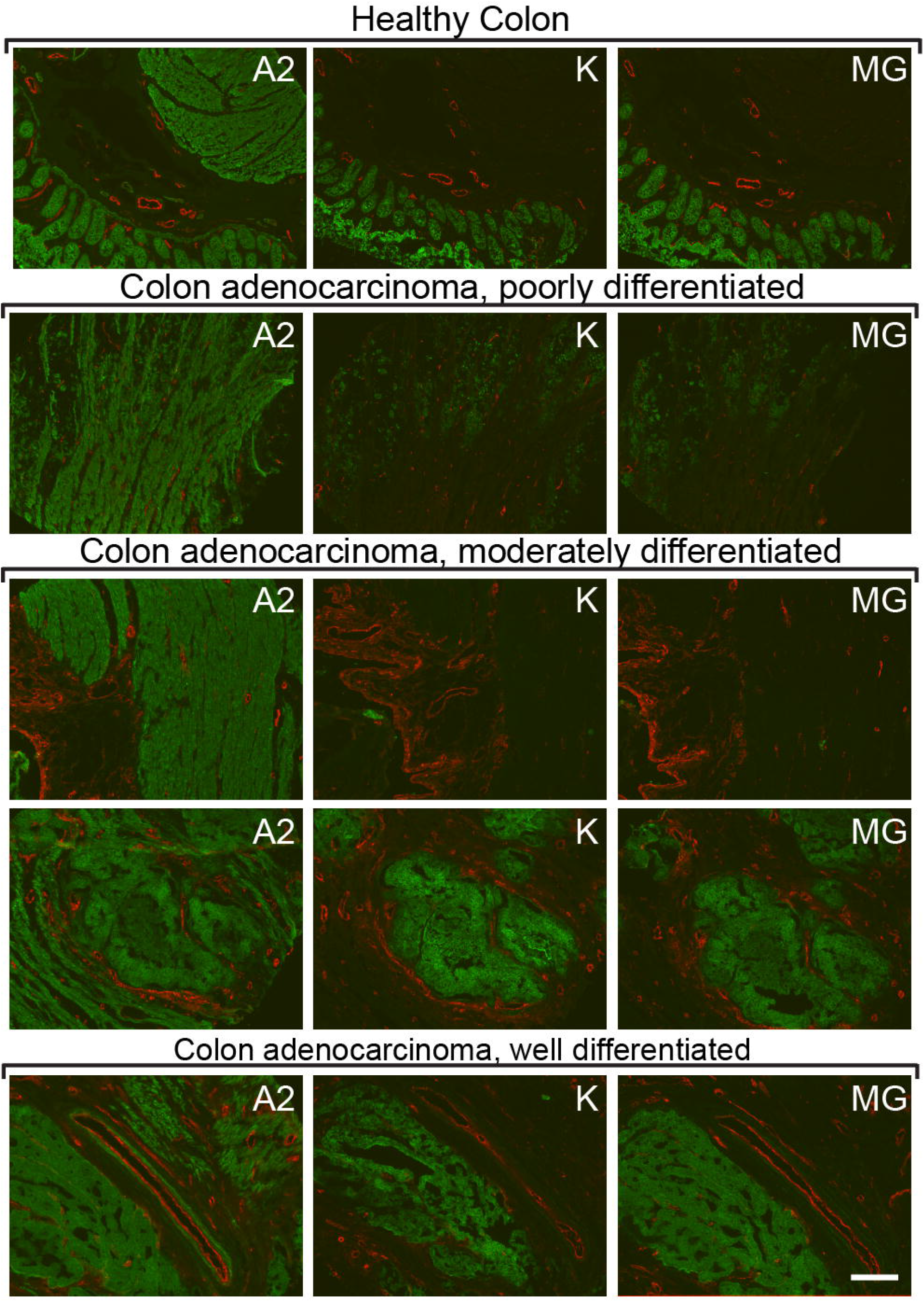
Immunofluorescence staining of A33 on human colon and human adenocarcinoma sections. Detection of A33 on human colon tissue sections and on human colon adenocarcinoma sections with the antibodies A2, K and MG (green, Alexa fluor 488). Blood vessels stained with an anti-Von Willebrand Factor antibody (red, Alexa fluor 595). Each horizontal line represents sequential sections from the same tumor sample, stained with a different antibody. Tumor samples vary in degree of differentiation. Scale bar = 150µm.

### Characterization of binding epitopes by SPOT technology

In order to identify the binding site of the anti-A33 antibodies, we used SPOT technology and immunodetection of antibody binding events to 15 amino acid long peptides on a cellulose membrane, spanning the antigen sequence. All three antibodies (A2, K and MG) recognized an immunodominant site of the A33 antigen [Figure 3b,c]. At a four times higher concentration, K and MG antibodies (2 µg/ml) generated multiple signals on the A33 peptide membrane, possibly indicating discontinuous binding epitopes on the Ig-like V-type (red) and Ig-like C2-type (green) domains [Figure 3b,c]. The TA99 antibody, serving as negative control of irrelevant specificity in this setting, did not exhibit binging on the SPOT assay [Figure 3c].

### Comparison of A33 detection on human tissues

The A2, K and MG antibodies were studied by immunofluorescence staining on a tissue array, which contained 37 colorectal cancer sections and 3 normal colon sections. The three antibodies were used at identical concentrations for the immunofluorescence analysis, while the TA99 antibody was used as negative control [**Supplementary Figure 3**]. Figure 4 shows sections stained in green with individual anti-A33 antibodies, while the vasculature was stained in red with an anti-Von Willebrand Factor reagent. The A2 antibody exhibited a brighter and more homogenous staining of tissue sections, compared to the K and MG antibodies, regardless of the tumor differentiation status [Figure 4]. A complete analysis of the tissue arrays can be found in **Supplementary Figure 3**.

### *In vivo* biodistribution of anti-A33 antibodies

The tumor-homing properties of the anti-A33 antibodies were assessed in immunocompetent BALB/c mice, bearing subcutaneously-grafted CT26^A33.C3^ carcinomas. An *ex vivo* immunofluorescence analysis, performed 24 hours after intravenous administration of A2, K and MG respectively, revealed a selective homing of the antibodies to the tumor cells [Figure 5]. By contrast, the negative control antibody TA99 did not exhibit any detectable accumulation at the tumor site at the same time point. A2 exhibited a more homogeneous and less patchy distribution within the tumor mass, compared to the K and MG counterparts.

**Figure 5.**
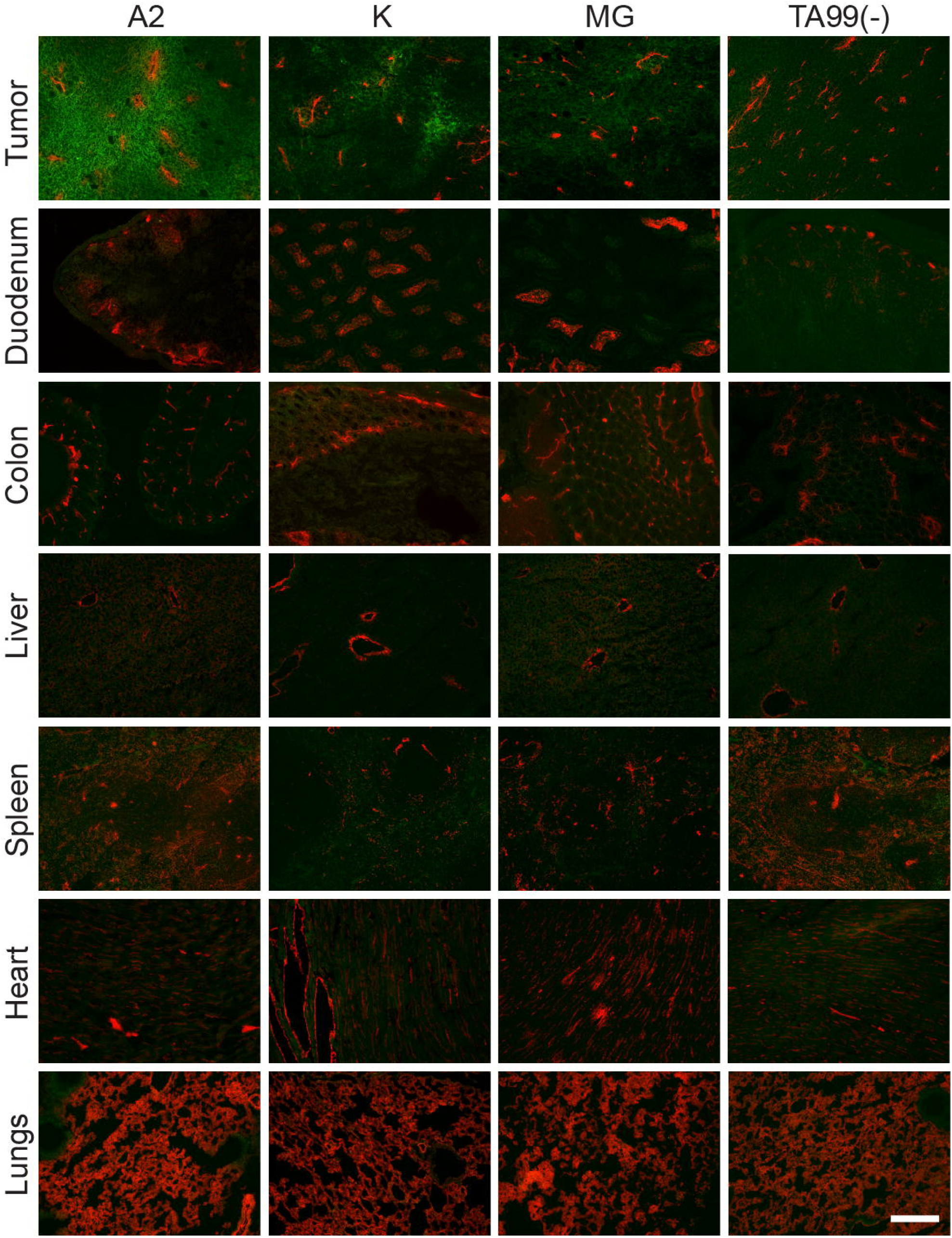
*In vivo* A2, K and MG selectively accumulate in s.c. CT26^A33.C3^ tumors. Microscopic fluorescence analysis of organs from CT26^A33.C3^ tumor bearing mice, 24 hours after intravenous administration of FITC labeled A2, K, MG or TA99 antibodies (green, AlexaFluor 488). Blood vessels stained with anti-CD31 (red, AlexaFluor 594). Magnification 10x, Scale bar = 125 µm.

### *In vitro* and *in vivo* A2-mediated cytotoxicity against C51 ^A33.C3^ and CT26^A33.C3^ colorectal cell lines

The anti-A33 antibody A2 was reformatted into human IgG1, in order to test its ability to induce ADCC *in vitro*, using human peripheral blood mononuclear cells (PBMCs). The sequence of IgG1(A2) and the protein characterization is reported in **Supplementary Figure 4**. The target cell lines C51^A33.C3^ and CT26^A33.C3^ were stained with 5(6)-Carboxyfluorescein diacetate N-succinimidyl ester (CFSE) dye and incubated with different concentrations of IgG1(A2) and human PMBCs, at 1:50 ratio. IgG1(KSF), directed against hen egg lysozyme was used as negative control in this setting. Flow cytometric analyses after 24 hours incubation revealed a specific and concentration-dependent depletion of CFSE-labeled target cells by the IgG1(A2) antibody [Figure 6a]. *In vivo*, the A2 antibody was used in the chimeric murine IgG2a format, in a setting aimed at preventing lung metastases. IgG2a(A2) was administered immediately after the intravenous injection of C51^A33.C3^ or CT26^A33.C3^ murine adenocarcinoma cells into BALB/c mice. The treatment significantly inhibited lung metastasis formation in both CT26 and C51 models, compared to saline or isotype-matched negative control antibodies (p < 0.05 for C51^A33.C3^ against isotype control and saline, p < 0.01 for CT26^A33.C3^ against isotype control and p < 0.001 against saline) [Figure 6b].

**Figure 6.**
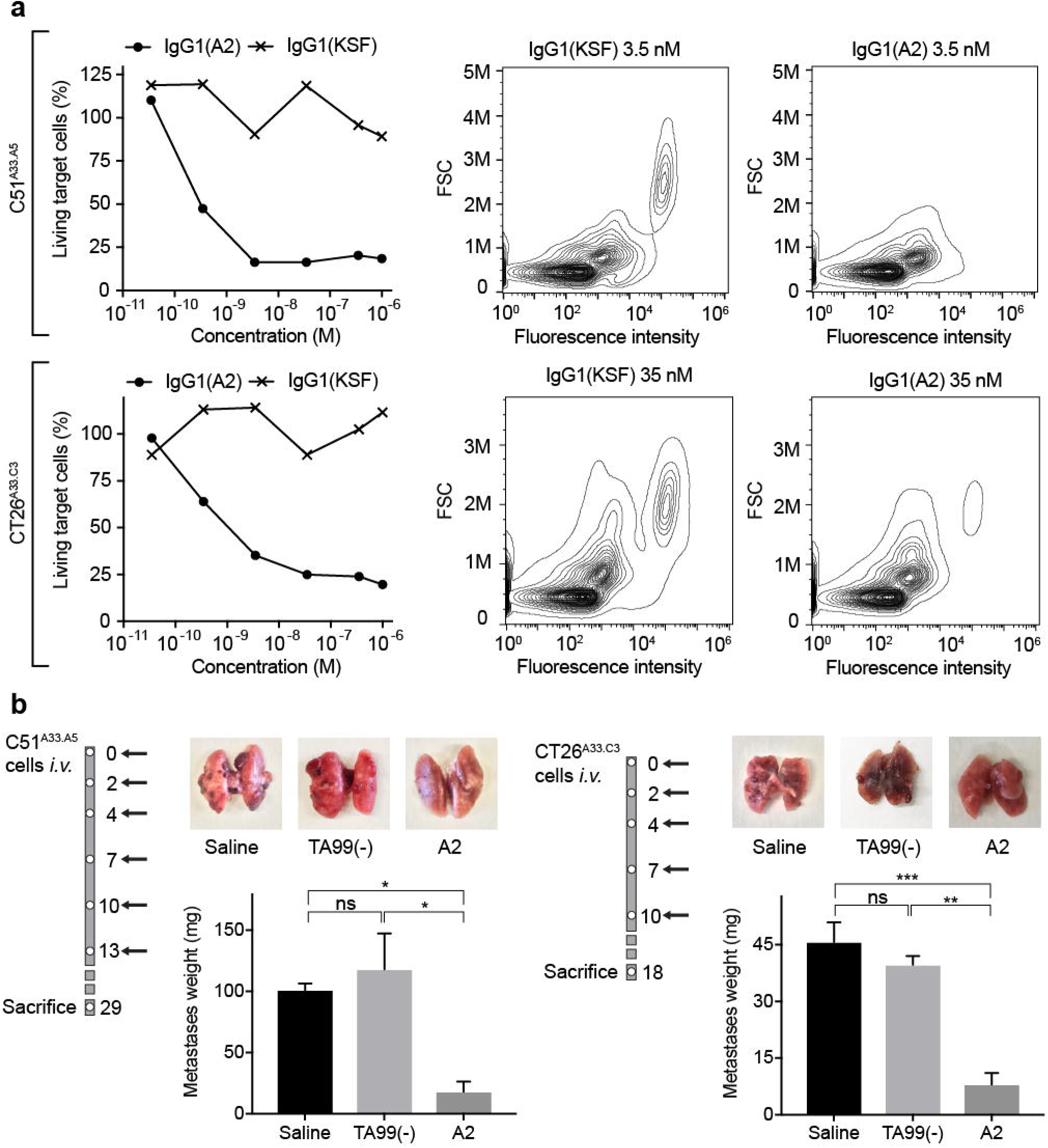
The antibody A2 induces ADCC *in vitro* and *in vivo.* **a)** *In vitro* ADCC killing of C51^A33.A5^ and CT26^A33.C3^ murine adenocarcinoma cells induced by the antibody A2 in IgG1 format, when incubated with human PBMCs for 24h. **Left:** Percentage of living tumor (CFSE positive) cells plotted against increasing antibody concentrations. **Right:** Representative FACS panels showing the specific lysis of tumor cells (CFSE stained). **b)** Inhibition of lung metastases formation upon treatment with the antibody A2 in the IgG2a format in mice injected intravenously with C51^A33.A5^ (left) CT26^A33.C3^ (right). Schematic representation of the therapy schedule indicating i.v. tumor cell injection, followed by 200µg A2, TA99 or saline i.v. injections. Representative images of lungs for each treatment group. Metastases were excised and weighted. Data represent mean metastases weight in mg ± SEM. ***, p < 0.001; **, p < 0.01;*, p < 0.05; ns, p > 0.05 (two-way ANOVA test, followed by Bonferroni posttest). C51^A33.A5^: Saline n = 3, TA99(-) and A2 n = 4. CT26^A33.C3^ n = 3.

## DISCUSSION

We have reported the generation of a novel fully-human antibody specific to the human tumor-associated antigen A33. The antibody, named A2, was compared *in vitro* and *in vivo* with two previously described anti-A33 antibodies, called K [17] and MG [9, 16].

The murine antibody K was tested in the early 90’s in Phase I/II clinical trials, in which the product was used as “delivery vehicle” for ^131^I [23] or ^125^I [24] in patients with metastatic CRC. A humanized version of the K antibody (referred to as K.hu), with longer serum half-life, was also used for radioimmunotherapy in radioiodinated form. The product did not induce objective tumor responses I patients, but a high accumulation in CRC metastases could be detected using Nuclear Medicine techniques [25]. Gastrointestinal toxicity was not reported, in spite of the fact that labeled K.hu antibody was found to also localize to the normal gastrointestinal tract [26]. After repeated monthly administrations, the humanized K.hu antibody induced a human anti-human antibody (HAHA) response in the majority of CRC patients (73%) [15], as evidenced by BIAcore analysis of sera. These data suggest that a less immunogenic anti-A33 antibody may be required for future clinical projects.

The MG antibody, generated by MacroGenics, has recently entered Phase I/II clinical trials in a bispecific antibody format [A33 x CD3 DART^™^ (MGD007)], aimed at recruiting T-cells to the tumor site. The product is also used in combination with an anti-PD-1 monoclonal antibody for the treatment of patients with metastatic CRC (NCT03531632). The bispecific T-cell engager MGD007 induced tumor growth inhibition in mouse models of colorectal cancer (LS174T and Colo205) (LS174T and Colo205) [16], but clinical results have not yet been disclosed.

The novel fully-human anti-A33 antibody A2 selectively and homogenously accumulated into subcutaneously-grafted CT26^A33.C3^ tumors in BALB/c mice upon i.v. injection [Figure 5]. Immunofluorescence staining of human colon adenocarcinoma and normal human colon tissue sections with A2 showed the presence of A33 in areas that resulted negative for A33 when stained with the antibodies K and MG [Figure 4]. The three antibodies showed to bind on an immunodominant epitope on the Ig-like V-type domain of A33 [Figure 3], even though the antibodies K and MG displayed binding on a discontinuous epitope. Binding of A2 to A33-expressing murine adenocarcinoma cells induced immune-mediated cell killing through ADCC as demonstrated with human PBMCs *in vitro* [Figure 6a] and by inhibition of lung metastases formation *in vivo* [Figure 6b].

ADCC represents an elegant avenue for re-directing the cytotoxic potential of immune effector cells to malignant cells, expressing a target antigen. We and others have previously shown that the main limitation for an efficient implementation of ADCC in a solid tumor setting relates to the lack of active immune effector cells in the neoplastic mass [27]. Clinical applications of monoclonal antibodies inducing ADCC for the treatment of disseminated solid tumors have been unsuccessful in most cases [28, 29]. However, the use of Trastuzumab as adjuvant treatment in metastatic breast cancer was shown to provide a substantial benefit to patients [30], suggesting that ADCC strategies may be better suited for the treatment of minimal residual disease. The good tolerability of intact antibodies make them suitable for adjuvant therapy, when long-term treatment is required. Alternatively, ADCC activity in solid tumors may be boosted by combination therapies with tumor-targeted immunostimulatory cytokines [27, 31, 32].

A33 expression in human CRC has been described to be heterogeneous [33, 34], as also observed by immunofluorescent staining of human CRC tissue sample [**Supplementary Figure 3**]. Therapeutic strategies capable of inducing a bystander effect on antigen-negative cells [35–37] may be particularly suited when targeting the A33 antigen.

The novel anti-A33 A2 antibody was able to successfully deliver payloads to the tumor site and to inhibit cancer dissemination in a minimal residual disease setting. A2 in IgG format may therefore be considered for adjuvant treatment applications in CRC patients or as a versatile “vehicle” for the delivery of bioactive payloads.

## MATERIALS AND METHODS

### Cell lines

CHO cells, the murine colorectal carcinoma cell line CT26^wt^ and the human colorectal cell lines HT-29 and LS174T were obtained from the ATCC between 2015 and 2017, expanded, and stored as cryopreserved aliquots in liquid nitrogen. Cells were grown according to the manufacturer’s protocol. Transfected CT26 cells were kept in the same culture conditions as CT26.wt cells.

### Cloning, expression, biotinylation and characterization of the antigen

The cDNA encoding for the A33 construct was obtained by total RNA extraction from the HT-29 colorectal carcinoma cell line (High Pure RNA Isolation Kit, Roche), reverse transcription (Transcriptor reverse transcriptase, Roche) and amplification of A33 cDNA by Taq DNA Polymerase (Sigma-Aldrich). The extracellular domains of the antigen were PCR amplified, extended with a nucleic acid sequence encoding for 6 histidine residues (His-tag) and cloned into the mammalian expression vector pcDNA3.1(+) (Invitrogen). The soluble antigen was produced by transient gene expression in CHO cells as described previously [38] and purified from the cell culture medium by Ni-NTA resin (Roche). Quality control of A33-His was performed by SDS-PAGE and size exclusion chromatography (Superdex75 10/300GL, GE Healthcare). The protein was digested with PNGase F (New England BioLabs) under denaturating conditions. A33-His was biotinylated with Sulfo-NHS-LC-Biotin (Pierce). Biotinylation of the antigen was tested through Mass Spectrometry (MS) analysis and band shift assay [**Supplementary Figure 5**].

### Transfection of human A33 in CT26 tumor cells and monoclonal selection

The gene for human A33 was cloned into the mammalian expression vector pcDNA3.1(+) containing an antibiotic resistance for G418 Geneticin. The plasmid was digested with PvuI (HF) for linearization prior to transfection. CT26^wt^ and C51^wt^ cells were transfected with the A33 gene in pcDNA3.1(+) using the Amaxa™ 4D-Nucleofector (Lonza) and the SG Cell Line 4D-Nucleofector® X Kit L (Lonza) and reseeded. Three days after the transfection, the growing medium was supplemented with 0.5 mg/ml G418 (Merck) for selection of stably transfected polyclonal cells. To obtain a monoclonal cell line, single cell sorting of stable transfected cells was performed using BD FACSAria III. Different clones were expanded and checked for antigen expression by FACS analysis and immunofluorescence. Clones CT26^A33.C3^ and C51^A33.A5^ were selected as the best clone based on A33 surface expression and cell viability.

### Selection of A33.A2 from the ETH-2-Gold library by phage display

The recombinant biotinylated A33-His antigen was immobilized on streptavidin (StreptaWells, Roche) or avidin coated wells (Avidin, Sigma-Aldrich on MaxiSorp plates, Sigma). Monoclonal antibodies were isolated from the synthetic human scFv antibody library ETH-2-Gold [19] by two rounds of biopanning as previously described [39, 40]. Supernatants of single bacterial colonies expressing the selected scFvs were screened by ELISA on immobilized A33-His. Clones yielding positive ELISA signals were sequenced and binding to the cognate antigen was confirmed by FACS analysis on the human colorectal carcinoma cell line LS174T and the transfected murine colorectal carcinoma cell line CT26^A33.C3^.

### Cloning, expression and characterization of the anti-A33 antibodies

Sequences of the variable region of the light and heavy chain (V_H_ and V_L_) of A33.K and A33.K.hu were taken from King D.J., et. al [18]. Sequences of A33.MG V_H_ and V_L_ were found in the patent file from MacroGenics [9]. The genes encoding for V_H_ and V_L_ of the antibodies A2, K, K.hu and MG, and the constant regions of the IgG2a immunoglobulin were PCR amplified, PCR assembled, and cloned into the mammalian expression vector pMM137, as previously described [27]. The IgG2a molecules were produced using transient gene expression in CHO cells following standard protocols [22, 38] and purified from the cell culture medium to homogeneity by Protein A chromatography (Thermo Scientific). Quality control of the proteins was performed by SDS-PAGE and size exclusion chromatography (Superdex200 10/300GL, GE Healthcare). The full amino acid sequences are presented in **Supplementary Figure 2**. The A2 antibody was additionally expressed in the human IgG1 format, for testing ADCC activity with human PBMCs. The sequence and characterization of IgG1(A2) are reported in **Supplementary Figure 4**.

### Surface plasmon resonance analysis on A33-his coated chip

Biotinylated A33-His was immobilized on SA-sensor chip with a density of 1000 RU using a BIAcore 200 System. Real-time interaction analysis was performed with serial dilutions of the anti-A33 antibodies. PBS (pH 7.4) was used as mobile phase and 10mM HCl as regeneration solution.

### Flow cytometry

CT26^wt^, CT26^A33.C3^, C51^wt^, C51^A33.A5^ and LS174T cells were detached with 2mM EDTA from the culture flask and stained with the different concentrations (75-0.001µg/ml) of the respective anti-A33 antibody or the negative control TA99. The antibodies were detected using donkey anti-mouse Alexafluor488 (Invitrogen). All staining and washing steps were performed in 2 mM EDTA 0.5% BSA in PBS. Cells were sorted by FACS (CytoFLEX, Beckman Coulter) and analyzed using FlowJo software.

### Epitope mapping using peptide array

The binding epitope of the anti-A33 antibodies on the A33 extracellular domain was determined using a peptide array (PepSpot, JPT). Sequences of 15 amino acid long peptides covering the whole sequence of the A33 extracellular domain with overlapping linear sequences were covalently bound to a cellulose membrane on 51 different spots. The assay was performed according to manufacturer’s instructions and the binding spots were detected by a chemiluminescence imager (Agfa Curix 60, Agfa Healthcare). The respective IgG2a antibodies were incubated with the PepSpot membrane at concentrations ranging from 0.5 µg/ml to 2µg/ml and detected by protein A-HRP (GE Healthcare).

### Ex-vivo immunofluorescences on CT26^A33.C3^ tumor bearing mice

7 weeks old BALB/c mice were obtained from Janvier. 4×10^6^. CT26^A33.C3^ cells were injected subcutaneously (s.c.) in the left flank of each mouse. The anti-A33 antibodies A2, K and MG were labelled with fluorescein isothiocyanate (FITC) as described in the manufacturer’s protocol (Sigma). For *ex vivo* immunofluorescence analysis, BALB/c mice bearing CT26^A33.C3^ s.c. tumors (100mm^3^) were injected with 150µg FITC-labelled IgG2a antibody. Mice were sacrificed 24 hours after injection. The organs were excised, embedded in cryoembedding medium (Thermo Scientific) and cryostat sections (8 mm) were stained using rabbit anti-FITC (BioRad; 4510-780) and rat anti-CD31 (BD Biosciences; 553370) as primary antibody and donkey anti-rabbit Alexa488 (Invitrogen; A21206) and anti-rat Alexa594 (Invitrogen; A21209) as secondary antibodies. Slides were mounted with fluorescent mounting medium (Dako) and analyzed with Axioscop2 mot plus microscope (Zeiss). *In vivo* experiments were performed under project licenses issued by the Veterinäramt des Kantons Zürich, Switzerland (Bew. Nr. 04/18).

### Immunofluorescence analysis on human colorectal tumor sections

Arrays of freshly frozen human colorectal tumor (37) and healthy colon tissues (3) were obtained from Amsbio. A list of the 40 different tissues, including pathology grade is shown in **Supplementary Figure 3**. For immunofluorescence analysis the tissue arrays were fixed in chilled acetone before staining. As primary staining antibodies A2, K, MG and TA99 were used at a concentration of 1µg/ml together with a rabbit anti-human Von Willebrand Factor (1:800, Dako) in 3% BSA in PBS. Detection was performed using a donkey anti-mouse IgG Alexafluor 488 (Invitrogen, A21202) and goat anti-rabbit IgG (Invitrogen, A11037).

### *In vitro* ADCC assay on CT26^A33.C3^ and C51^A33.A5^ cells

CT26^A33.C3^ and C51^A33.A5^ were stained with CFSE dye (BioLegend) as described by the manufacturer’s protocol and seeded in a 48-well plate (2·10^4^/well). Peripheral blood samples were obtained from healthy donors from the Blutspendedienst SRK, Zurich. PBMCs were isolated by density gradient centrifugation on Ficoll Paque Plus (GE Healthcare) following the manufacturer’s protocol. The peripheral blood was diluted 1:3 in 2 mM EDTA / PBS. 30 ml of diluted peripheral blood were layered on 12.9 ml Ficoll and centrifuged at 400g for 40 min at RT. PBMCs were collected, washed twice with PBS/EDTA and incubated with the tumor target cells (10^6^/well, 1:50 target:effector ratio) in CT26^A33.C3^ and C51^A33.A5^ culture media. The antibody IgG1(A2) was added at different concentrations to the wells (total volume of 200µl/well) and the plate was incubated at 37°C, 5% CO_2_ for 24 hours. On the next day, the samples with IgG1(A2) + PBMCs + target cells were transferred to a 96-well plate, washed twice with PBS and stained Fixable Viability Dye (Invitrogen) for 30 min. The samples were washed twice with 2 mM EDTA 0.5% BSA in PBS, before being analyzed by FACS (CytoFLEX, Beckman Coulter). The percentage of living target cells was derived from the mean fluorescence intensity of the CFSE positive cell population normalized based on the negative control samples containing PBMCs and target cells only.

### *In vivo* prevention of CT26^A33.C3^ and C51^A33.A5^ lung metastases formation

5·10^5^ CT26^A33.C3^ or 5·10^4^ C51^A33.A5^ cells were injected intravenously through the tail vein of 7 weeks old BALB/c mice (Janvier). Mice were injected with saline, 200µg of A2 or TA99 antibody in the IgG2a format as indicated in **Figure 7b** (black arrows). After euthanasia, mice were perfused with 1% PFA through the pulmonary artery and the aorta after. Lungs were excised and fixed for 1h in 4% PFA. Lung metastases were excised and weighted. Unresectable, visible metastases were counted and added to the final weight (1mg each). Experiments were performed under project licenses issued by the Veterinäramt des Kantons Zürich, Switzerland (Bew. Nr. 04/18).

## ACKNOWLEDGEMENTS

We thank Dr. Renier Myburgh for providing the PBMCs for *in vitro* ADCC evaluation. The authors gratefully acknowledge financial support from ETH Zürich, the ERC Advanced Grant “Zauberkugel” (Grant Agreement 670603), the Swiss National Science Foundation (project number: 310030_182003/1), the Swiss Federal Commission for Technology and Innovation (Grant number: 17072.1) and the “Stiftung zur Krebsbekämpfung”.

## ABBREVIATIONS

ADCC: Antibody-dependent cell-mediated cytotoxicity
CDR: Complementarity-determining region
CEA: Carcinoembyogenic antigen
CFSE: 5(6)-Carboxyfluorescein diacetate N-succinimidyl ester
CRC: Colorectal cancer
EGFR: Endothelial growth factor receptor
HAHA: Human anti-human antibody
Mw: Molecular weight
PBMC: Peripheral blood mononuclear cell
scFv: Single chain variable fragment
VEGF: Vascular endothelial growth factor
V_L_: Variable domain of the light chain
V_H_: Variable domain of the heavy chain

## ETHICS APPROVAL AND CONSENT TO PARTICIPATE

Experiments were performed under a project license (license number 04/2018), granted by the Veterinäramt des Kanton Zürich, Switzerland, in compliance with the Swiss Animal Protection Act (TSchG) and the Swiss Animal Protection Ordinance (TSchV).

## REFERENCES

1. Bray F, Ferlay J, Soerjomataram I, Siegel RL, Torre LA, Jemal A. Global cancer statistics 2018: GLOBOCAN estimates of incidence and mortality worldwide for 36 cancers in 185 countries. CA Cancer J Clin. 2018; 68: 394–424. doi: 10.3322/caac.21492.

2. Riihimaki M, Hemminki A, Sundquist J, Hemminki K. Patterns of metastasis in colon and rectal cancer. Sci Rep. 2016; 6: 29765. doi: 10.1038/srep29765.

3. Marley AR, Nan H. Epidemiology of colorectal cancer. Int J Mol Epidemiol Genet. 2016; 7: 105–14. doi:

4. Kalyan A, Kircher S, Shah H, Mulcahy M, Benson A. Updates on immunotherapy for colorectal cancer. J Gastrointest Oncol. 2018; 9: 160–9. doi: 10.21037/jgo.2018.01.17.

5. Overman MJ, Lonardi S, Wong KYM, Lenz HJ, Gelsomino F, Aglietta M, Morse MA, Van Cutsem E, McDermott R, Hill A, Sawyer MB, Hendlisz A, Neyns B, et al. Durable Clinical Benefit With Nivolumab Plus Ipilimumab in DNA Mismatch Repair-Deficient/Microsatellite Instability-High Metastatic Colorectal Cancer. J Clin Oncol. 2018; 36: 773–9. doi: 10.1200/JCO.2017.76.9901.

6. Zanzonico P, Carrasquillo JA, Pandit-Taskar N, O’Donoghue JA, Humm JL, Smith-Jones P, Ruan S, Divgi C, Scott AM, Kemeny NE, Fong Y, Wong D, Scheinberg D, et al. PET-based compartmental modeling of (124)I-A33 antibody: quantitative characterization of patient-specific tumor targeting in colorectal cancer. Eur J Nucl Med Mol Imaging. 2015; 42: 1700–6. doi: 10.1007/s00259-015-3061-2.

7. Schwegler C, Dorn-Beineke A, Nittka S, Stocking C, Neumaier M. Monoclonal anti-idiotype antibody 6G6.C4 fused to GM-CSF is capable of breaking tolerance to carcinoembryonic antigen (CEA) in CEA-transgenic mice. Cancer Res. 2005; 65: 1925-33. doi: 10.1158/0008-5472.CAN-04-3591.

8. Klein C, Waldhauer I, Nicolini VG, Freimoser-Grundschober A, Nayak T, Vugts DJ, Dunn C, Bolijn M, Benz J, Stihle M, Lang S, Roemmele M, Hofer T, et al. Cergutuzumab amunaleukin (CEA-IL2v), a CEA-targeted IL-2 variant-based immunocytokine for combination cancer immunotherapy: Overcoming limitations of aldesleukin and conventional IL-2-based immunocytokines. Oncoimmunology. 2017; 6: e1277306. doi: 10.1080/2162402X.2016.1277306.

9. Moore PA, Li J, Zhifen Chen F, Johnson LS, Shah K, Bonvini E. (2015). Bi-specific diabodies that are capable of binding gpA33 and CD3 and uses thereof MacroGenics Inc).

10. Behr T, Becker W, Hannappel E, Goldenberg DM, Wolf F. Targeting of liver metastases of colorectal cancer with IgG, F(ab’)2, and Fab’ anti-carcinoembryonic antigen antibodies labeled with 99mTc: the role of metabolism and kinetics. Cancer Res. 1995; 55: 5777s-85s. doi:

11. Goldenberg DM, Wlodkowski TJ, Sharkey RM, Silberstein EB, Serafini AN, Garty, II, Van Heertum RL, Higginbotham-Ford EA, Kotler JA, Balasubramanian N, et al. Colorectal cancer imaging with iodine-123-labeled CEA monoclonal antibody fragments. J Nucl Med. 1993; 34: 61–70. doi:

12. Murray JL, Rosenblum MG, Zhang HZ, Podoloff DA, Kasi LP, Curley SA, Chan JC, Roh M, Hohn DC, Brewer H, et al. Comparative tumor localization of whole immunoglobulin G anticarcinoembryonic antigen monoclonal antibodies IMMU-4 and IMMU-4 F(ab’)2 in colorectal cancer patients. Cancer. 1994; 73: 850–7. doi:

13. Carrasquillo JA, Pandit-Taskar N, O’Donoghue JA, Humm JL, Zanzonico P, Smith-Jones PM, Divgi CR, Pryma DA, Ruan S, Kemeny NE, Fong Y, Wong D, Jaggi JS, et al. (124)I-huA33 antibody PET of colorectal cancer. J Nucl Med. 2011; 52: 1173–80. doi: 10.2967/jnumed.110.086165.

14. Garinchesa P, Sakamoto J, Welt S, Real F, Rettig W, Old L. Organ-specific expression of the colon cancer antigen A33, a cell surface target for antibody-based therapy. Int J Oncol. 1996; 9: 465–71. doi:

15. Welt S, Ritter G, Williams C, Jr., Cohen LS, John M, Jungbluth A, Richards EA, Old LJ, Kemeny NE. Phase I study of anticolon cancer humanized antibody A33. Clin Cancer Res. 2003; 9: 1338–46. doi:

16. Moore PA, Shah K, Yang Y, Alderson R, Roberts P, Long V, Liu D, Li JC, Burke S, Ciccarone V, Li H, Fieger CB, Hooley J, et al. Development of MGD007, a gpA33 x CD3-Bispecific DART Protein for T-Cell Immunotherapy of Metastatic Colorectal Cancer. Mol Cancer Ther. 2018; 17: 1761–72. doi: 10.1158/1535-7163.MCT-17-1086.

17. Welt S, Divgi CR, Real FX, Yeh SD, Garin-Chesa P, Finstad CL, Sakamoto J, Cohen A, Sigurdson ER, Kemeny N, et al. Quantitative analysis of antibody localization in human metastatic colon cancer: a phase I study of monoclonal antibody A33. J Clin Oncol. 1990; 8: 1894–906. doi: 10.1200/JCO.1990.8.11.1894.

18. King DJ, Antoniw P, Owens RJ, Adair JR, Haines AM, Farnsworth AP, Finney H, Lawson AD, Lyons A, Baker TS, et al. Preparation and preclinical evaluation of humanised A33 immunoconjugates for radioimmunotherapy. Br J Cancer. 1995; 72: 1364–72. doi: 10.1038/bjc.1995.516.

19. Silacci M, Brack S, Schirru G, Marlind J, Ettorre A, Merlo A, Viti F, Neri D. Design, construction, and characterization of a large synthetic human antibody phage display library. Proteomics. 2005; 5: 2340–50. doi: 10.1002/pmic.200401273.

20. Welt S, Mattes MJ, Grando R, Thomson TM, Leonard RW, Zanzonico PB, Bigler RE, Yeh S, Oettgen HF, Old LJ. Monoclonal antibody to an intracellular antigen images human melanoma transplants in nu/nu mice. Proc Natl Acad Sci U S A. 1987; 84: 4200–4. doi: 10.1073/pnas.84.12.4200.

21. Frank R. The SPOT-synthesis technique. Synthetic peptide arrays on membrane supports--principles and applications. J Immunol Methods. 2002; 267: 13–26. doi:

22. Pasche N, Woytschak J, Wulhfard S, Villa A, Frey K, Neri D. Cloning and characterization of novel tumor-targeting immunocytokines based on murine IL7. J Biotechnol. 2011; 154: 84–92. doi: 10.1016/j.jbiotec.2011.04.003.

23. Welt S, Divgi CR, Kemeny N, Finn RD, Scott AM, Graham M, Germain JS, Richards EC, Larson SM, Oettgen HF, et al. Phase I/II study of iodine 131-labeled monoclonal antibody A33 in patients with advanced colon cancer. J Clin Oncol. 1994; 12: 1561–71. doi: 10.1200/JCO.1994.12.8.1561.

24. Welt S, Scott AM, Divgi CR, Kemeny NE, Finn RD, Daghighian F, Germain JS, Richards EC, Larson SM, Old LJ. Phase I/II study of iodine 125-labeled monoclonal antibody A33 in patients with advanced colon cancer. J Clin Oncol. 1996; 14: 1787–97. doi: 10.1200/JCO.1996.14.6.1787.

25. Chong G, Lee FT, Hopkins W, Tebbutt N, Cebon JS, Mountain AJ, Chappell B, Papenfuss A, Schleyer P, U P, Murphy R, Wirth V, Smyth FE, et al. Phase I trial of 131I-huA33 in patients with advanced colorectal carcinoma. Clin Cancer Res. 2005; 11: 4818–26. doi: 10.1158/1078-0432.CCR-04-2330.

26. Scott AM, Lee FT, Jones R, Hopkins W, MacGregor D, Cebon JS, Hannah A, Chong G, U P, Papenfuss A, Rigopoulos A, Sturrock S, Murphy R, et al. A phase I trial of humanized monoclonal antibody A33 in patients with colorectal carcinoma: biodistribution, pharmacokinetics, and quantitative tumor uptake. Clin Cancer Res. 2005; 11: 4810–7. doi: 10.1158/1078-0432.CCR-04-2329.

27. Murer P, Kiefer JD, Pluss L, Matasci M, Blumich SL, Stringhini M, Neri D. Targeted Delivery of TNF Potentiates the Antibody-Dependent Cell-Mediated Cytotoxicity of an Anti-Melanoma Immunoglobulin. J Invest Dermatol. 2018. doi: 10.1016/j.jid.2018.11.028.

28. Slamon DJ, Leyland-Jones B, Shak S, Fuchs H, Paton V, Bajamonde A, Fleming T, Eiermann W, Wolter J, Pegram M, Baselga J, Norton L. Use of chemotherapy plus a monoclonal antibody against HER2 for metastatic breast cancer that overexpresses HER2. N Engl J Med. 2001; 344: 783–92. doi: 10.1056/NEJM200103153441101.

29. Van Cutsem E, Kohne CH, Hitre E, Zaluski J, Chang Chien CR, Makhson A, D’Haens G, Pinter T, Lim R, Bodoky G, Roh JK, Folprecht G, Ruff P, et al. Cetuximab and chemotherapy as initial treatment for metastatic colorectal cancer. N Engl J Med. 2009; 360: 1408–17. doi: 10.1056/NEJMoa0805019.

30. Baselga J, Perez EA, Pienkowski T, Bell R. Adjuvant trastuzumab: a milestone in the treatment of HER-2-positive early breast cancer. Oncologist. 2006; 11 Suppl 1: 4–12. doi: 10.1634/theoncologist.11-90001-4.

31. Schliemann C, Palumbo A, Zuberbuhler K, Villa A, Kaspar M, Trachsel E, Klapper W, Menssen HD, Neri D. Complete eradication of human B-cell lymphoma xenografts using rituximab in combination with the immunocytokine L19-IL2. Blood. 2009; 113: 2275–83. doi: 10.1182/blood-2008-05-160747.

32. Gutbrodt KL, Schliemann C, Giovannoni L, Frey K, Pabst T, Klapper W, Berdel WE, Neri D. Antibody-based delivery of interleukin-2 to neovasculature has potent activity against acute myeloid leukemia. Sci Transl Med. 2013; 5: 201ra118. doi: 10.1126/scitranslmed.3006221.

33. Baptistella AR, Salles Dias MV, Aguiar S, Jr., Begnami MD, Martins VR. Heterogeneous expression of A33 in colorectal cancer: possible explanation for A33 antibody treatment failure. Anticancer Drugs. 2016; 27: 734–7. doi: 10.1097/CAD.0000000000000379.

34. Uhlen M, Fagerberg L, Hallstrom BM, Lindskog C, Oksvold P, Mardinoglu A, Sivertsson A, Kampf C, Sjostedt E, Asplund A, Olsson I, Edlund K, Lundberg E, et al. Proteomics. Tissue-based map of the human proteome. Science. 2015; 347: 1260419. doi: 10.1126/science.1260419.

35. Bremer E, Samplonius D, Kroesen BJ, van Genne L, de Leij L, Helfrich W. Exceptionally potent anti-tumor bystander activity of an scFv:sTRAIL fusion protein with specificity for EGP2 toward target antigen-negative tumor cells. Neoplasia. 2004; 6: 636–45. doi: 10.1593/neo.04229.

36. Kovtun YV, Audette CA, Ye Y, Xie H, Ruberti MF, Phinney SJ, Leece BA, Chittenden T, Blattler WA, Goldmacher VS. Antibody-drug conjugates designed to eradicate tumors with homogeneous and heterogeneous expression of the target antigen. Cancer Res. 2006; 66: 3214–21. doi: 10.1158/0008-5472.CAN-05-3973.

37. Ross SL, Sherman M, McElroy PL, Lofgren JA, Moody G, Baeuerle PA, Coxon A, Arvedson T. Bispecific T cell engager (BiTE(R)) antibody constructs can mediate bystander tumor cell killing. PLoS One. 2017; 12: e0183390. doi: 10.1371/journal.pone.0183390.

38. Rajendra Y, Kiseljak D, Baldi L, Hacker DL, Wurm FM. A simple high-yielding process for transient gene expression in CHO cells. J Biotechnol. 2011; 153: 22–6. doi: 10.1016/j.jbiotec.2011.03.001.

39. Ravenni N, Weber M, Neri D. A human monoclonal antibody specific to placental alkaline phosphatase, a marker of ovarian cancer. MAbs. 2014; 6: 86–94. doi: 10.4161/mabs.27230.

40. Viti F, Nilsson F, Demartis S, Huber A, Neri D. Design and use of phage display libraries for the selection of antibodies and enzymes. Methods Enzymol. 2000; 326: 480–505. doi:

